# Proof-of-Concept in a Murine Model of Treatment of Thrombotic Thrombocytopenic Purpura Using Engineered Red Blood Cells

**DOI:** 10.1101/2025.06.12.659386

**Authors:** Karl S Roberts, Shouping Zhang, Khulan Batbayar, Zi Yan, Joshua Muia, J Justin Mulvey, Emmanuel Olivier, James M Pullman, Spero R Cataland, Eric E Bouhassira

## Abstract

Thrombotic Thrombocytopenic Purpura (TTP) is caused by congenital or acquired deficiency of ADAMTS13, a metalloproteinase that cleaves von Willebrand Factor (vWF) multimers. Current treatments—plasma exchange and immunosuppression—are costly and associated with significant morbidity therefore, alternative strategies are needed.

We developed the kitJak2 platform for producing genetically engineered lab-grown red blood cells (lgRBCs) as drug delivery vectors. We hypothesized that membrane-bound ADAMTS13 displayed on lgRBCs could provide a durable treatment for TTP. To test this, we engineered erythroid cells expressing both wild-type and mutant variants MDTCS fragments of ADAMTS13, conferring resistance to autoantibodies. Flow cytometry and FRET-based assays confirmed robust membrane expression and enzymatic activity. Importantly, mutant MDTCS variants retained catalytic activity in the presence of plasma from TTP patients, whereas wild-type variants were inhibited.

For in vivo evaluation, we generated transgenic mice expressing MDTCS ADAMTS13 on their RBC membranes. These mice exhibited normal RBC half-lives and stable, catalytically active ADAMTS13 expression. Using a murine model of TTP—where ADAMTS13 knockout mice challenged with recombinant human vWF (rhvWF) develop thrombocytopenia and schistocytes—we demonstrated that transfusion of ADAMTS13-expressing RBCs significantly mitigated disease, preventing platelet loss and schistocyte formation. This confirms that membrane-bound MDTCS ADAMTS13 cleaves circulating rhvWF under physiological flow conditions in vivo.

Finally, employing our KitJak2 platform, we generated human enucleated lgRBCs expressing high levels of catalytically active ADAMTS13.

This novel work establishes proof-of-concept that membrane-anchored ADAMTS13-expressing lab-grown RBCs may offer a feasible and effective therapeutic approach for both congenital and acquired TTP.

## Introduction

Thrombotic Thrombocytopenic Purpura (TTP) is a life-threatening disorder caused by a deficiency of A Disintegrin and Metalloproteinase with Thrombospondin Type 1 Motif, Member 13 (ADAMTS13).^1^ This deficiency can be either congenital (cTTP) or, more commonly, due to acquired inhibitory autoantibodies in immune-mediated TTP (iTTP). ADAMTS13 prevents excessive platelet aggregation by cleaving multimers of the pro-coagulant glycoprotein von Willebrand Factor (vWF) conformationally altered by high levels of shear stress in the circulation. Insufficient ADAMTS13 activity results in the accumulation of shear-stress activated vWF multimers, leading to widespread platelet aggregation and formation of microthrombi in arterioles where shear is at its highest level.

The standard treatment for TTP has been plasma infusion for cTTP and daily plasma exchange combined with immunosuppressive therapy for iTTP.^2,3^ Recently, two novel therapies have received FDA approval. Caplacizumab, a humanized, bivalent nanobody that inhibits the interaction between vWF and platelets was approved to prevent microvascular thrombosis in iTTP, while Adzynma a recombinant ADAMTS13 (rADAMTS13) was approved as an enzyme replacement therapy to restore ADAMTS13 activity in cTTP. Adzyma is also currently undergoing clinical trial for iTTP.

These two therapies, potentially in combination with plasma exchange, may become the preferred first-line therapy for TTP, pending the outcomes of ongoing clinical trials.^4^ Nevertheless, despite these advances, TTP is increasingly recognized as a chronic, relapsing disease. Relapses are common, and many patients who survive the acute phase experience incomplete recovery and long-term complications, including cerebrovascular and cardiovascular events.^5,6^ The underlying mechanisms driving these chronic sequelae are not fully understood but may be linked to persistent fluctuations or deficiencies in ADAMTS13 activity.

Regular infusions of recombinant rADAMTS13, combined with routine monitoring of ADAMTS13 activity,^7^ may offer an effective prophylactic strategy. However, the development of improved methods for sustained, long-term delivery of ADAMTS13 remains a significant unmet need.

Genetically modified stem cell-derived lab-grown red blood cells (lgRBCs) have long been explored as a potential platform for drug delivery, owing to their natural 120-day lifespan.^8^ Several protocols have been developed to differentiate and expand primary hematopoietic stem and progenitor cells (HSPCs) or immortalized erythroid cells and differentiate them into enucleated red blood cells. ^9–18^ These methods hold a lot of promises but suffer from significant drawbacks, including reliance on primary cells or expensive culture conditions, karyotypic instability,^19^ and efficient enucleation only at low cell densities, all of which hinder their suitability for large-scale clinical application.

To overcome these challenges, we have recently developed a strategy that addresses many of the key limitations. This approach is centered on the discovery that self-renewing erythroblasts (SREs) can be consistently derived from induced pluripotent stem cells (iPSCs) engineered to carry the kitD816V and jak2V617F mutations.^20^ These KitJak2 SREs exhibit karyotypic stability and demonstrate robust expansion capacity without the need for cytokines, in a chemically defined, albumin-free, cost-effective medium.^21^ Crucially, KitJak2 SREs retain the ability to differentiate efficiently and enucleate into mature lgRBCs at high rates. These properties make KitJak2 SREs an ideal, scalable source for producing genetically engineered lgRBCs suitable for drug delivery applications In this study, we demonstrate in a mouse model that transfusions of ADAMTS13-expressing RBCs effectively prevent the onset of TTP. Furthermore, we show that human lgRBCs derived from KitJak2 SREs can be genetically modified to express high levels of ADAMTS13, providing a scalable platform for therapeutic applications.

## Results

### Catalytically Active ADAMTS13 Fragments Can Be Tethered to the Cell Membrane

To assess the feasibility of expressing ADAMTS13 on the surface of erythroid cells, we initially employed K562 cells due to their ease of genetic manipulation and robust activity of erythroid-specific promoters. Given the large size of full-length ADAMTS13, we focused on its N-terminal 650 amino acid fragment, known as the MTDCS fragment, which comprises the metalloprotease domain, thrombospondin type 1 repeat, disintegrin-like domain, the cysteine-rich and the spacer domain (**Fig. 1A**). This region retains catalytic activity comparable to the full-length protein while lacking most of the epitopes targeted by autoantibodies in iTTP.^22^

**Figure 1:**
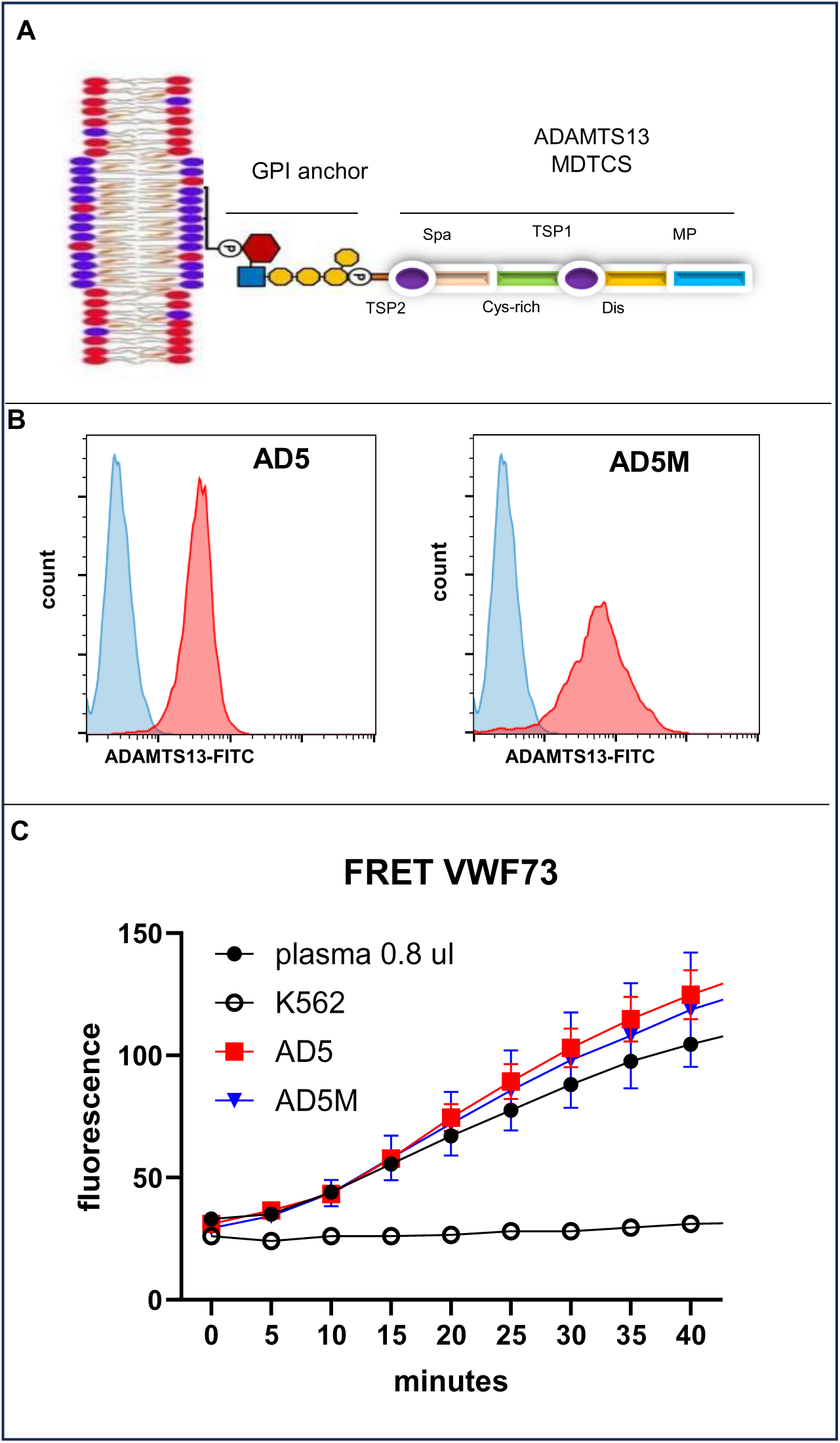
GPI-anchored ADAMTS13 expression in K562 cells. A: Schematic representation of the GPI-anchored, membrane-bound MTDCS fragment of ADAMTS13, illustrating its structure following membrane insertion. **B:** Surface expression of ADAMTS13. Constructs encoding the MTDCS ADAMTS13 fragment fused to a GPI-anchored membrane molecule, and driven by globin gene regulatory elements were knocked into the AAVS1 sites in K562 cells. Expression of both the wild-type (AD5) and antibody-resistant (AD5M) fusion proteins was confirmed by flow cytometry, with both variants expressed at high levels. **C:** FRET analysis demonstrated that the membrane-bound MTDCS fragments retain catalytic activity. 15,000 K562 cells were compared to 0.8uL of human plasma (n=3).

Since ADAMTS13 is naturally secreted, we designed a membrane-anchored construct termed AD5, containing the full MTDCS fragment fused to a glycophosphatidylinositol (GPI) anchor sequence derived from Decay Acceleration Factor (DAF), as previously described.^23^This GPI anchor facilitates integration into the plasma membrane via lipid rafts during protein synthesis.

In addition to AD5, we generated four other fusion constructs: AD5M which encode a gain-of-function MTDCS variant resistant to most of the auto-antibodies that causes iTTP (**Fig. S1**),^24^ and three smaller sub-fragments of MTDCS. All constructs were placed under the control of an erythroid-specific α-globin promoter (HBA2) coupled with elements of the β-globin locus control region to ensure high-level expression in erythroid cells.^25^ Following plasmid assembly, the fusion constructs were stably integrated into the AAVS1 safe harbor locus on chromosome 19 of K562 cells using CRISPR/Cas9 technology.^25^ Flow cytometry with an antibody targeting the N-terminal region of ADAMTS13 confirmed surface expression of the constructs (**Fig. 1B**).

To evaluate catalytic activity, we performed a FRET-based assay using a 73-amino acid fragment of vWF containing a fluorophore-quencher pair flanking the cleavage site, enabling direct measurement of proteolytic activity.^26^ Citrated plasma and EDTA plasma were used as positive and negative controls, respectively. Both AD5 and AD5M constructs demonstrated robust catalytic activity, with 15,000 K562 cells displaying equivalent activity to 1 µL of human plasma (**Fig. 1C**). In contrast, smaller MTDCS sub-fragments showed no detectable activity. Based on these results, we selected the AD5 and AD5M constructs for subsequent experiments.

### Generation and Characterization of ADAMTS13-containing RBCs in mice

Having demonstrated that it is possible to generate cells expressing a catalytically active ADAMTS13 fragment, we next produced transgenic mice expressing ADAMTS13 on their RBCs to assess whether this expression would impact their half-life. To achieve this, we inserted an AD5 construct into the Rosa26 locus. Although founder mice were successfully generated, flow cytometry analysis revealed heterogeneous expression, even after breeding to homozygosity (**Fig. 2A**). To achieve more uniform expression levels, we subsequently used pronuclear injections to create transgenic mice harboring multiple copies of the AD5M construct inserted at random genomic loci. Multiple founder lines were obtained; however, expression remained heterogeneous across all lines. A founder with AD5 inserted at rosa26 and an AD5M founder obtained by pronuclear injections were selected and bred to homozygosity for use in subsequent experiments (**Fig. 2A**).

**Figure 2:**
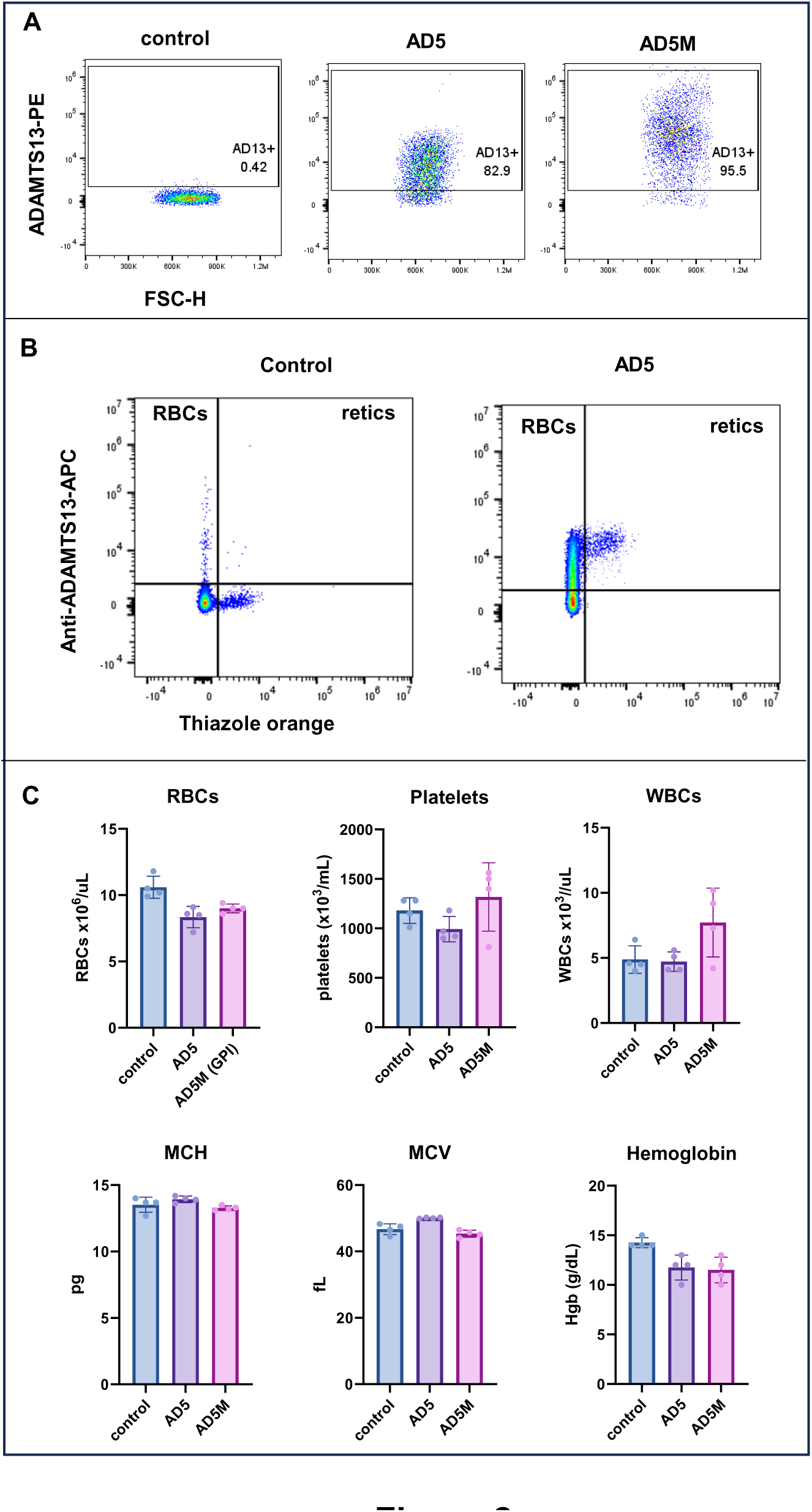
GPI-anchored ADAMTS13 expression in mouse RBCs. **A:** Constructs encoding the MTDCS fragment of ADAMTS13 fused to a GPI-anchored membrane molecule, driven by erythroid-specific regulatory elements, were introduced into the Rosa26 locus in mice using CRISPR/Cas9 technology (AD5 line) or by pronuclear injection into mouse oocytes (AD5M line). Surface expression of the MTDCS fragment on RBC membranes was confirmed by FACS analysis. While the MTDCS fragment was readily detectable, expression was heterogeneous, even when the transgenes were bred to homozygosity. **B:** Mouse washed RBCs were co-labeled with thiazole orange (to identify reticulocytes) and anti- ADAMTS13-APC antibodies. Reticulocytes uniformly exhibited high levels of ADAMTS13 expression, whereas mature RBCs displayed a heterogeneous expression pattern. This suggests robust expression of the constructs in erythroid precursors, with progressive loss of surface expression as RBCs age in circulation. **C:** Blood samples from control mice and from mice expressing the AD5 and AD5M constructs were analyzed using an ADVIA 2120i automated hematology analyzer. No significant differences were observed in complete blood counts or red cell indices among the three groups (n=4).

To investigate the source of the heterogeneous expression, we co-stained RBCs with anti-ADAMTS13 antibodies and thiazole orange to identify reticulocytes. This analysis revealed that reticulocytes consistently displayed high levels of membrane-bound ADAMTS13 fragments (**Fig. 2B**), indicating an inverse correlation between fusion protein expression and cell age—reticulocytes being high expressers and mature erythrocytes exhibiting lower levels. These findings suggest that the constructs are robustly expressed in reticulocytes as they exit the bone marrow, but the membrane-bound protein is progressively lost as the reticulocytes mature into erythrocytes and age in circulation.

Mice expressing ADAMTS13 fragments appeared healthy and bred normally. To specifically evaluate whether expression of ADAMTS13 fragments impacts RBC viability or function, we first analyzed RBC counts and hematological parameters in the AD5 and AD5M mice. The analysis showed no significant differences in either RBC indices or total cell numbers between transgenic and control mice, indicating that expression of ADAMTS13 fragments does not lead to anemia (**Fig. 2C**).

To confirm and extend these findings, we performed a half-life assay, comparing transgenic cells to wild-type mouse RBCs. Cells from both groups were labeled with one of two distinct lipophilic fluorescent dyes, DiI or DiD, mixed in equal proportions, and transfused into recipient mice. Blood samples were collected at multiple time points over a 30-day period, and fluorescence signal decay was monitored to assess RBC survival. Both wild-type and ADAMTS13-expressing RBCs exhibited similar rates of signal loss, indicating comparable lifespan in vivo (**Figs. 3A & B**). However, consistent with our earlier observations, although the overall half-life of the transgenic RBCs was unaffected, the expression of the ADAMTS13 fragment progressively declined over time in DiD-labeled cells (**Fig. S2**). This confirms that while expression is robust initially, the membrane-bound GPI-anchored fusion protein is not stably maintained in circulating mouse RBCs.

**Figure 3:**
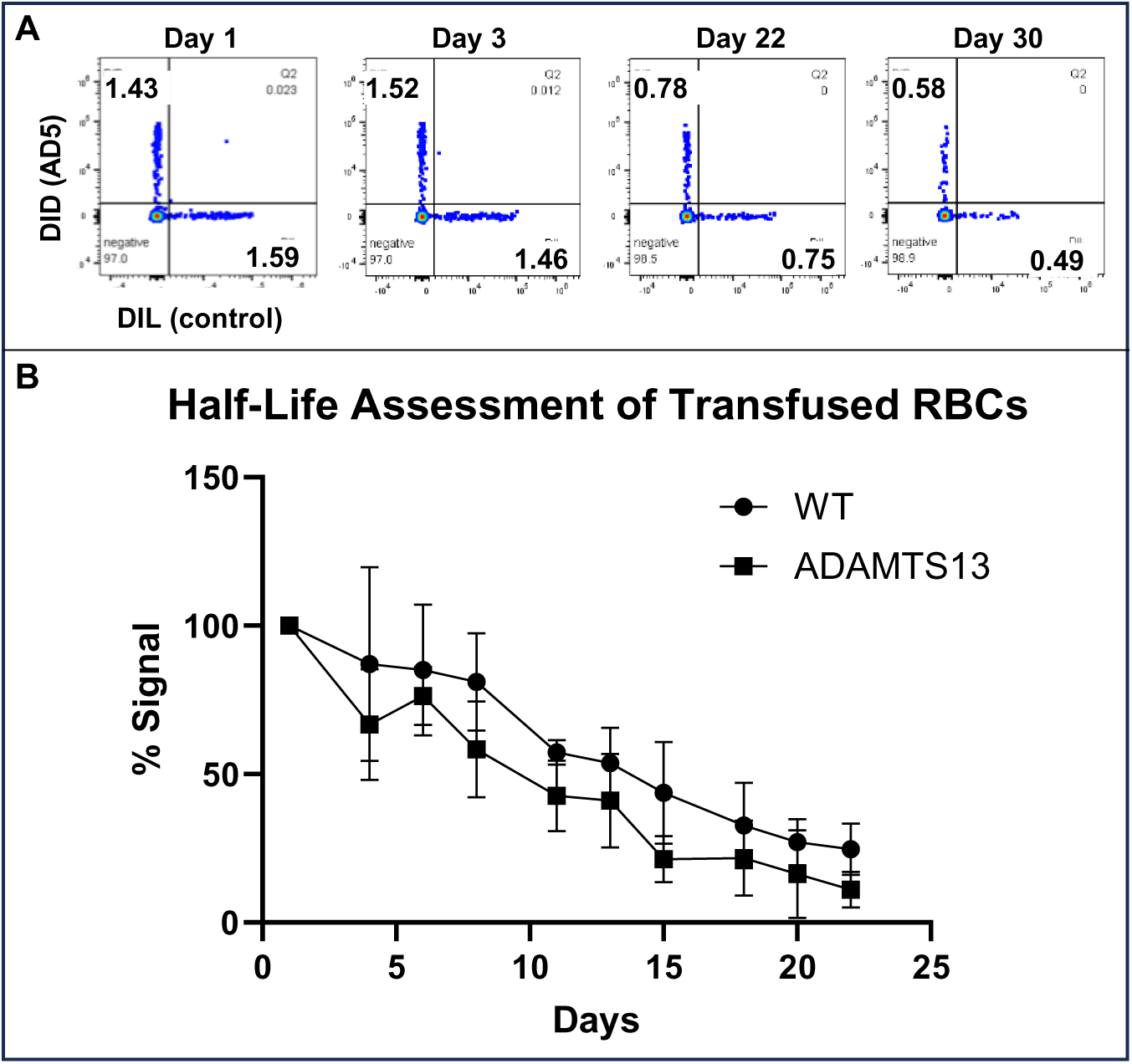
GPI-anchored ADAMTS13 does not alter RBC lifespan in vivo. **A:** Approximately 200 million control and AD5-RBCs were labeled with either DID or DIL lipophilic dyes and co-injected intraorbitally into control mice (8 to 12 weeks old). RBC survival was monitored over time by FACS analysis. **B:** The percentage of labeled RBCs relative to the initial time point (1hour post-injection) is plotted (n=4). Similar results were observed with AD5M cells. The decay curves for control and AD5 RBCs show no significant differences, indicating that expression of membrane-bound ADAMTS13 does not alter RBC lifespan in vivo.

### Proof-of-Concept: Transfusion of ADAMTS13-containing reticulocytes can reduce or prevent the onset of TTP symptoms

While deficiency of ADAMTS13 leads to congenital cTTP in humans, expression of an overt, acute cTTP event, often requires a triggering event such as infection, pregnancy or excessive alcohol use. Even in the case of iTTP, very low levels of ADAMTS13 are not always immediately associated with a crisis.

The ADAMTS13 knockout (ADAMTS13KO) mice have a phenotype similar to humans. They do not exhibit an overt phenotype, ^27,28^ but a murine model mimicking key features of TTP has been established by Shiviz et al. by injecting recombinant human vWF (rhvWF) into ADAMTS13KO mice.^29^ To assess whether AD5M-RBCs could prevent the onset of TTP symptoms in this model, we conducted a transfusion challenge in which ADAMTS13KO mice received either wild-type or ADAMTS13-expressing RBCs prior to rhvWF administration (**Fig. S3**).

Since reticulocytes exhibited higher levels of ADAMTS13 expression than RBCs—and because lgRBCs are in the reticulocyte stage when generated—we chose to perform these experiments using reticulocytes rather than mature RBCs. To produce the reticulocytes, donor AD5M mice were treated with 50 mg/kg phenyl-hydrazine, which selectively eliminates the existing RBC population and stimulates robust reticulocyte production.

FACS analysis performed 48 hours after phenylhydrazine treatment revealed a marked increase in the percentage of reticulocytes, rising from approximately 2% to over 80%. Consistently, FRET-based assays using both VWF73 and VWF71^30^—a FRET substrate specifically designed to minimize quenching by hemoglobin³⁰—demonstrated robust catalytic activity on the membranes of murine RBCs expressing AD5 or AD5M. Notably, this activity was more than three-fold higher in reticulocytes compared to mature RBCs. (**Figs. 4A & B**)

**Figure 4:**
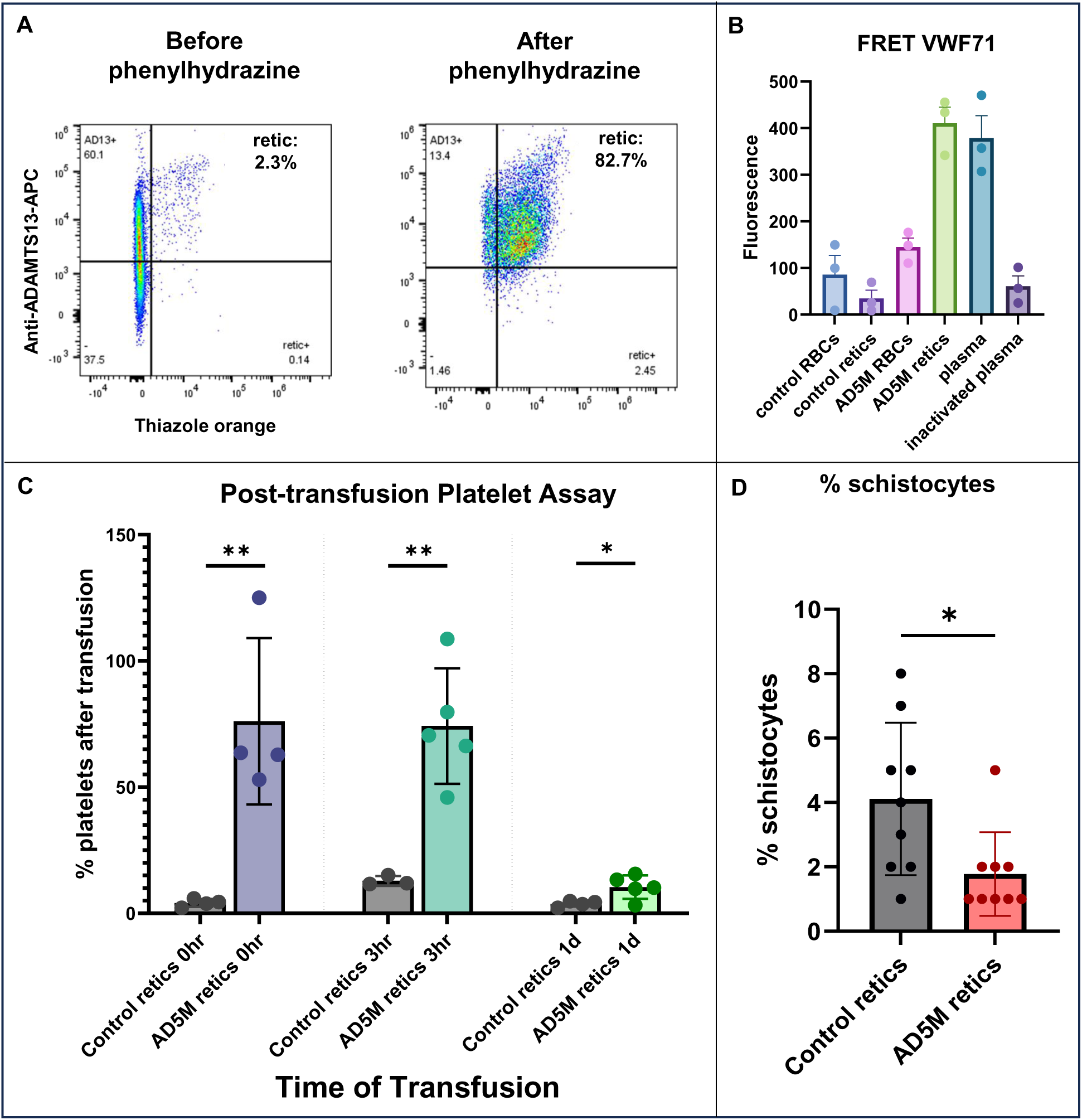
Proof-of-concept for TTP treatment using AD5M-RBCs. **A:** Washed peripheral RBCs were co-stained with thiazole orange and ADAMTS13-APC antibodies before and 48 hours after phenylhydrazine injection. FACS analysis shows a marked increase in reticulocyte percentage post-treatment, with virtually all reticulocytes expressing the AD5M construct. **B:** ADAMTS13 catalytic activity was assessed in RBCs and reticulocytes collected before and after phenylhydrazine treatment, respectively, using the FRET VWF71 assay. Activity was compared to human plasma (1.2 μL) and inactivated plasma (EDTA-treated). Bar graph demonstrates significantly higher activity in reticulocytes (n=4). **C:** ADAMTS13 knockout mice were transfused with approximately 200 million AD5M-RBCs and challenged with 2000 IU/kg rhVWF at 0-, 3-, or 24-hours post-transfusion. Platelet counts were measured using an ADVIA 2120i automated hematology analyzer. (n=4 to 5 depending on time points). The plots show that AD5M-RBC transfusion fully or partially prevents the platelet drop typically observed in this TTP model. **D:** Mice were transfused with AD5M-RBCs and challenged with vWF at 0 or 3 hours post-transfusion. Peripheral blood smears were stained with Wright-Giemsa stain, and schistocyte counts were assessed by microscopy 24 hours post-rhvWF. The plot shows reduced schistocyte formation in transfused mice (n=8).

Forty-eight hours after phenyl-hydrazine administration, approximately 200 million AD5M reticulocytes were harvested and transfused by retro-orbital injection into ADAMTS13KO mice. Shortly after the transfusion the mice were challenges with 2,000 IU/kg of rhvWF to induce TTP-like symptoms.^29^ Blood samples were then acquired at periodic interval in order to measure the platelet and the schistocyte counts.

Following administration of rhVWF, ADAMTS13KO mice, whether transfused with wild-type reticulocytes or not, exhibited a near-complete depletion of circulating platelets within one day. In contrast, mice transfused with reticulocytes expressing membrane-bound ADAMTS13 (AD5M-retics) at the time of rhVWF injection showed minimal to no reduction in platelet counts (**Fig. 4C**). These results suggest that membrane-anchored ADAMTS13 might cleaves full-length exogenous rhVWF, thereby preventing pathological platelet consumption. Furthermore, transfusion of AD5-retics up to 24 hours prior to rhVWF administration blocked or significantly mitigated platelet loss and resulted in fewer schistocytes compared to controls (**Figs. 4C & D**). Collectively, these findings highlight that transfusion of AD5M-retics can substantially reduce or prevent key hematological features of TTP in ADAMTS13KO mice.

The Schiviz TTP model is associated with cardiac and renal damage, which can be prevented or reduced by administration of ADAMTS13. To evaluate whether transfusion of AD5M-expressing reticulocytes could similarly protect against tissue injury, ADAMTS13KO mice were transfused with either control or AD5M reticulocytes, followed by injection of rhvWF. Organs were harvested three days later, fixed, embedded in paraffin, and sectioned for Hematoxylin and Eosin staining. The heart was the only organ to show damage and inflammation (**Fig. S4**), in the control mice. Inflammation was multifocal and was composed mostly of lymphocytes and granulation tissue in damaged cardiomyocytes. Control mouse kidney, pancreas and spleen showed no significant inflammation. All mice treated with AD5M- expression reticulocytes were free of inflammation and tissue injury in all organs examined. We conclude that reticulocytes expressing MTDCS fragments can completely prevent tissue damage, as evidenced by the absence of leukocyte infiltration in cardiac tissue. (**Fig. S4**)

### Specific fragments of ADAMTS13 exhibit resistance to anti-ADAMTS13 autoantibodies present in TTP patients

While the above experiments demonstrate that ADAMTS13 fragments can be successfully attached to the RBC membrane via a GPI -anchor, expression was heterogeneous and did not persist in aged RBCs. To improve the expression of membrane-bound ADAMST13 fragment, we elected to modify the fusion protein by swapping out the GPI anchor for either the full length or the transmembrane domain of Glycophorin B, a 91 amino acid transmembrane protein native to RBCs. We also generated an additional antibody-resistant constructs, containing a single point mutation (K608N) that has been shown to alter the N-glycosylation profile of the protein (**Fig. S2B**).^31^ Flow cytometry and FRET analysis of K562 cell lines generated from these new constructs show increased membranous expression and catalytic activity (**Figs. 5A & B**). Constructs containing the full length ADAMTS13 fused to the N-terminal of Glycophorin B (**Fig. S1**) were poorly expressed and were not pursued any further (data not shown). To test for antibody resistance in vitro, we incubated the transgenic K562 cell lines with plasma harvested from TTP patients and conducted a FRET assay using cells incubated with normal plasma as a negative control. Plotting of the residual catalytic activity after plasma incubation showed that K562 lines lacking the antibody resistant mutations were sensitive to all of the patient plasma samples as shown by a significant decrease in catalytic activity compared to controls (**Fig. 5C**). K562 cell lines with the gain-of-function AD5M mutations were resistant to 11 of the 13 patients plasma samples, while lines with the K608N-AD5 fragment were resistant to all of the plasma tested (13/13). This leads us to conclude that the antibody resistant mutations that we tested can confer resistance to anti-ADAMTS13 autoantibodies of iTTP patients in the context of membrane-bound MTDCS fragment.

**Figure 5:**
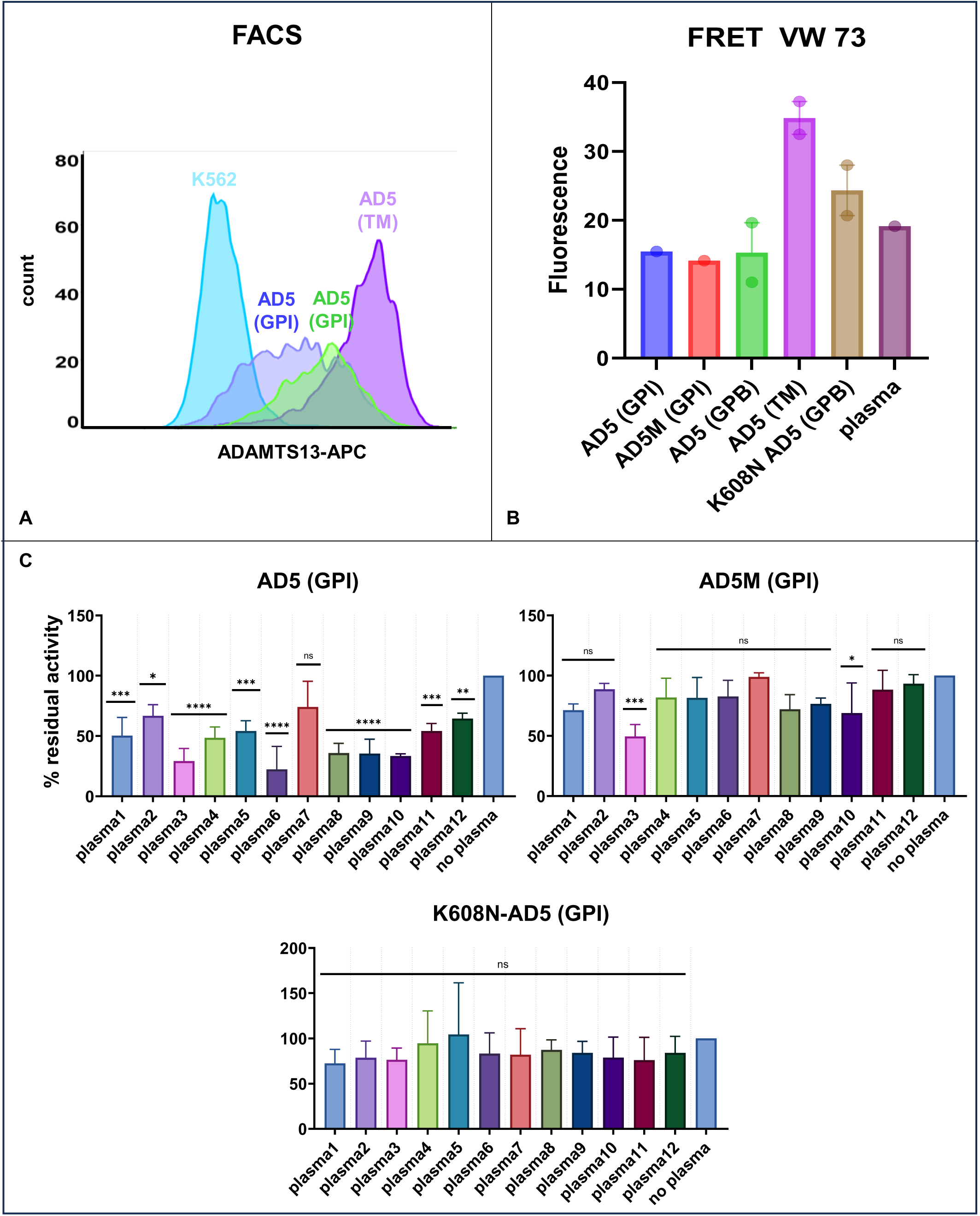
Gain-of-function and K608N mutation confer autoantibody resistance to membrane-bound ADAMTS13 fragments. **A:** To enhance membrane expression, the GPI anchor used in the AD5 construct was replaced with either the full-length GPB coding sequence or its transmembrane domain. The modified constructs were knocked into the AAVS1 locus of K562 cells. Flow cytometry histograms show that membrane expression is higher when the MTDCS fragment is fused to either the full-length GPB or its transmembrane domain compared to the original DAF GPI anchor. **B:** FRET analysis using the VWF73 assay confirms that the increased expression correlates with higher catalytic activity. **C:** K562 cells expressing AD5, AD5M, or K608N-AD5 fragments were incubated for 30 minutes with plasma from 13 TTP patients. After washing, residual ADAMTS13 activity was measured via the VWF73 FRET assay. The plot shows that AD5 activity is inhibited by nearly all patient plasma samples, whereas AD5M resists inhibition by 11/13 plasma samples, and the K608N-AD5 fragment maintains activity in all cases, indicating complete resistance to autoantibody inhibition.

### Membrane-bound MTDCS fragments cleave full-length high-molecular-weight vWF multimers

Building on the FRET and in vivo data presented above, which strongly suggested that membrane-bound MTDCS fragments can cleave full-length high-molecular-weight vWF multimers, we sought to directly confirm this activity. RhvWF was incubated for 1 hour at 37°C with control K562 cells or K562 cells expressing MTDCS fragments. Multimer analysis by agarose gel electrophoresis revealed that MTDCS fragments efficiently cleaved full-length vWF into monomers and smaller multimers. (**Fig. 6**)

**Figure 6:**
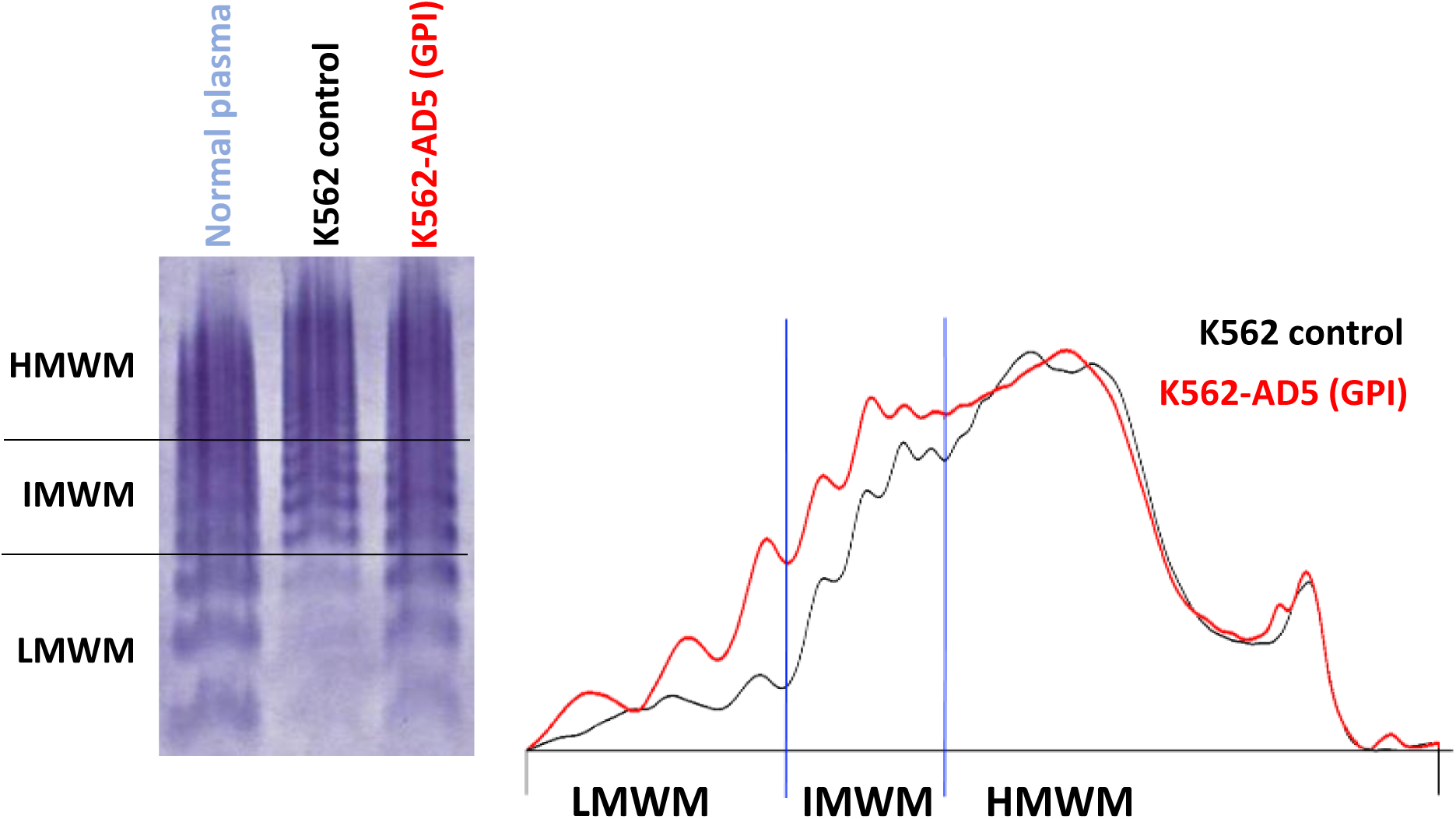
Membrane-bound MTDCS fragments cleave full-length high-molecular-weight vWF multimers. RhvWF was incubated for 1 hour at 37 °C with control K562 cells or K562 cells expressing MTDCS fragments. Representative micrograph of vWF multimer analysis by agarose gel electrophoresis demonstrates that MDTCS fragments efficiently cleaved full-length vWF into monomers and lower-order multimers.

### Production of human lab-grown RBCs that express membrane-bound ADAMTS13 fragments

To evaluate the feasibility of expressing antibody-resistant ADAMTS13 fragments on human RBCs, we inserted the K608N-AD5 construct into KitJak2 iPSCs at the AAVS1 safe harbor locus using CRISPR/Cas9 gene-editing technology (**Fig 7A)**. These modified iPSCs were then used to generate KitJak2 SREs, which were subsequently differentiated into lgRBCs, as previously described.²⁰ ³¹

**Figure 7:**
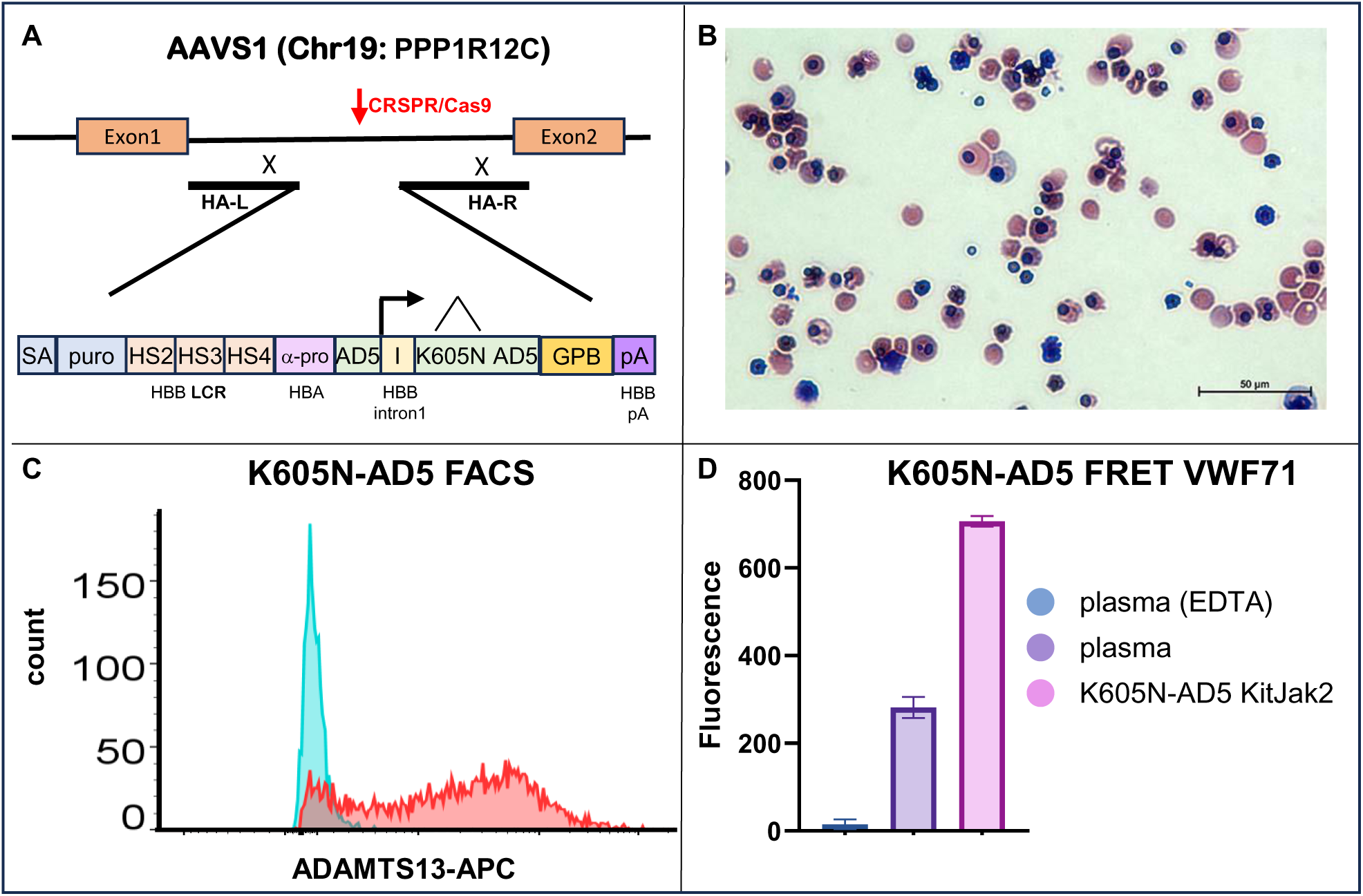
Expression of membrane-bound K608N-AD5 MTDCS fragment in lab-grown human RBCs. **A:** A construct encoding the K608N-AD5 fragment fused to full-length GPB was knocked into the AAVS1 site of KitJak2 iPSCs, and SREs were generated. **B:** The SREs were differentiated for 10 days in a bioreactor and analyzed by Giemsa staining. Approximately 50% of the resulting cells were enucleated. **C & D:** FACS analysis and FRET VWF71 assays demonstrated high-level surface expression and robust catalytic activity of the membrane-bound K608N-AD5 MTDCS fragment in the lab-grown RBCs. Catalytic activity of 150,000 cells was compared to 10 μL of plasma (n=2).

Flow cytometry analysis and Giemsa staining confirmed that expression of the K608N-AD5 fusion protein did not impair enucleation, and that the fusion protein was expressed at high levels (**Figs. 7B & C**). Functional analysis using FRET-based VWF70 assays revealed that approximately 50,000 K608N-AD5- expressing lgRBCs exhibited ADAMTS13 activity equivalent to 1 µL of plasma (**Fig. 7D**). Based on this activity, we estimate that a transfusion of approximately 0.17 mL of packed lgRBCs per kg body weight (equivalent to ∼12 mL for a 70 kg individual) could provide the same ADAMTS13 activity as a prophylactic dose of 40 IU/kg recombinant ADAMTS13 (rADAMTS13).

These results demonstrate that lgRBCs expressing high levels of the K608N-AD5 fusion protein can be efficiently generated using KitJak2 technology, supporting the feasibility of this approach for therapeutic application.

## Discussion

Recent advances in therapeutic strategies for TTP have significantly improved the management of acute iTTP, to the extent that the disease is increasingly evolving into a chronic condition. In part due to the novelty of these treatments, the underlying drivers of relapse and long-term complications remain incompletely understood. However, it is widely believed that sustained normalization of ADAMTS13 activity could alleviate many of these clinical challenges.

Several approaches exist for delivering therapeutic proteins to the plasma over a patient’s lifetime. The most straightforward strategy involves periodic infusion of the therapeutic protein, ideally modified to prolong its half-life. While this method is feasible, it poses potential drawbacks, including high costs, potential adverse effects, and the risk of poor patient adherence due to the need for frequent visits to healthcare facilities.

An alternative strategy for long-term therapy is gene therapy. In physiological conditions, ADAMTS13 is primarily produced by hepatic stellate cells. While genetically modifying these native cells to express therapeutic, antibody-resistant variants of ADAMTS13 would offer an elegant, targeted solution, the technology to achieve precise gene editing in stellate cells is currently not available.^32^

In this study, we demonstrate that expression of a fragment of ADAMTS13 anchored to the membrane of RBCs effectively prevents excessive platelet consumption and schistocyte formation—two hallmarks of TTP. Unlike stellate cells, RBCs are highly accessible to gene therapy approaches using lentiviral vectors. In fact, the constructs employed in this work closely resemble the lentiviral vectors that we and others developed over two decades ago for gene therapy targeting hemoglobinopathies.^33^. Notably, variations of these constructs have since gained FDA approval for clinical use in treating hemoglobinopathies.

Taken together, our findings suggest that gene therapy leading to stable expression of membrane-bound, antibody-resistant variants of ADAMTS13 in RBCs could offer a viable and durable therapeutic approach to prevent relapses and achieve long-term disease control in TTP following initial acute management.

Gene therapy using this approach, which involves an autologous stem cell transplant, remains a complex and costly intervention. Additionally, it carries inherent risks of treatment failure, and may therefore not be suitable for all TTP patients.

A third alternative to the sustained delivery of therapeutic protein involves the use of therapeutic transfusion of engineered lgRBCs. This approach has long been theorized but has so far never been reduced to practice, although a phase III clinical trial is in progress for a variation of this approach involving physical loading of small molecules into autologous RBCs^34^.

A key barrier has been the high cost and inefficiencies associated with the production of genetically modified RBCs, particularly challenges in scalability, stability, and genetic manipulation. Our recently developed technology addresses many of these limitations. Specifically, KitJak2 SREs are essentially immortal, karyotypically stable, highly amenable to genetic modification, and can be produced at a significantly reduced cost, providing a feasible and scalable platform for the generation of engineered lgRBCs for therapeutic use.

In this study, we demonstrate that our technology enables the efficient production of lgRBCs expressing high levels of the K608N-AD5 fragment. It is generally accepted that maintaining ADAMTS13 activity above 20% of normal is sufficient to prevent subclinical microangiopathic hemolysis and thrombocytopenia, which may contribute to the reduced life expectancy observed in iTTP survivors^5,6^. Depending on the half-life of K608N-AD5-expressing RBCs in human circulation, transfusion of small volumes—potentially just a few milliliters or sub-milliliter amounts—of packed RBCs could be enough to sustain protective ADAMTS13 levels for several months at a time.

Our technology involves introducing oncogenic mutations into iPSCs to generate stable RBC precursors (SREs). This is not considered a barrier to clinical application, as the resulting lgRBCs are enucleated and can be irradiated prior to transfusion to eliminate any residual nucleated cells. The iPSCs used to produce the current KitJak2 SRE line carry the blood group O, RhD-negative phenotype.

However, repeated transfusions—even of small volumes—might still pose a risk of alloimmunization if mismatches in minor blood group antigens exist. Further studies will be required to evaluate this risk. I In any cases, the risk could be significantly mitigated by matching, or genetically knocking-out, key immunogenic antigens such as Rh (CcEe), Kell, Duffy, and Kidd, or by generating autologous lgRBCs derived from the patient’s own cells.

Additionally, while the expression of fusion proteins and the incorporation of mutations conferring resistance to autoantibodies could theoretically provoke an immune response, the GPB-K608N-AD5 fusion protein used in this study is composed of two naturally occurring proteins linked together, and contains only a single amino acid substitution. This design minimizes the likelihood of immunogenicity.

In conclusion, we present a proof-of-concept demonstrating that lgRBCs expressing fragments of ADAMTS13, engineered to resist autoantibody inhibition, can effectively prevent key clinical features of TTP in an animal model. Furthermore, these therapeutic lgRBCs can be efficiently produced using our KitJak2 SRE technology. Such cells may prove valuable for the treatment of cTTP, acute iTTP, and for prophylaxis.

## Methods

### Internal Review Board

All experiments involving human cells were conducted under protocols approved by the Albert Einstein College of Medicine Institutional Review Board (IRB), Bronx, New York.

### Constructs

Plasmids used for the knock-in of ADAMTS13 fusion proteins were based on previously described backbones.^25^ All fusion constructs were generated using a combination of DNA synthesis and standard molecular biology techniques. Complete plasmid sequences are available upon request.

### CRISPR Knock-In at AAVS1

K562 cells were electroporated with plasmids encoding the synthetic guide RNA and Cas9 mRNA using a custom electroporation system. Puromycin selection was applied 48 hours post-electroporation to enrich for successfully edited cells. PCR analysis of the junction fragments on both sides of the insertion site confirmed that over 95% of the recovered clones carried the correct integration at the AAVS1 locus

### Cell culture

K562 were obtained from the ATCC and cultured as suggested by ATCC. iPSCs line O1 and O2 have been described previously ^21^.

### Self-renewing erythroblasts (SRE)

#### Differentiation of iPSCs into kitjak2 SREs

Day 0: kitjak2 iPSCs were differentiated into day-17 HPCs according to the previously published protocol.^20^ day-17 HPCs were then cultured in IMIT combined with 1mM Dex, 30mM IBMX. After 10 to 14 days, greater than 98% of the cells in the culture had acquired the antigen profiles of SREs.

#### Long-term culture of SREs

kitjak2 SREs can be cultured for about 120 days in IMIT combined with 1uM Dex and 30uM IBMX.^20^ All SREs were passaged every 3 to 5 days by dilution to 1.25 to 2.5x10^5^ cells/mL once the culture concentration exceeded 1.5x10^6^/mL.

#### Terminal differentiation of SREs

Day 0: Cells were centrifuged, rinse once in PBS and plated at about 1.5x10^5^ cells/mL in R6 media containing 4U/ml of Epo, 5% human plasma and 1mM RU 486. Day 3: Cells were diluted 1 to 2 to about 3.5x10^5^ cells/mL in the same media without Epo. Days 5, 7 and 9: Cells were diluted in pure RPMI to about 3 to 5x10^5^ cells/mL.

### Transfusion Challenge

ADAMTS13 knockout (ADAMTS13KO) mice were obtained from Jackson Laboratory and used as the recipient strain. C57BL/6 mice, aged 6 to 8 weeks, were transfused via retro-orbital injection with 50- 100ul of packed reticulocytes harvested from either wild-type donors or ADAMTS13-expressing mice. Following transfusion, ADAMTS13KO recipients were administered recombinant human von Willebrand factor (VONVENDI) at a dose of 2,000 U/kg body weight, delivered via the tail vein at 0, 3, and 24 hours post-transfusion. Peripheral blood was collected before and after VONVENDI administration, and complete blood counts, including platelet quantification, were performed using an ADVIA 2120i automated hematology analyzer.

### Hematoxylin and Eosin staining

Mice were euthanized day 4 post-transfusion in accordance with an IACUC-approved protocol. Kidney, heart, liver spleen and pancreas tissues were formalin fixed and submitted for routine histologic processing, paraffin embedding and microtomy to create 4 micron thick sections mounted on glass slides and stained with hematoxylin and eosin. Slides were examined with an Olympus BX73 brightfield microscope and photographed with a Nikon DS-Fi3 5.9 megapixel camera at objective magnifications of 4X and 40X.

### FRET VWF73 K562 Catalytic Assay

K562 cell lines expressing or lacking ADAMTS13 were washed, centrifuged, and resuspended in VWF73 assay buffer (Sensolyte) at a concentration of 50,000 cells/µL. For each condition, 10 µL of the cell suspension was dispensed into a 384-well plate, followed by the addition of 10 µL of fluorescently labeled VWF73 substrate (Sensolyte). Fluorescence was recorded every 2.5 minutes for 30 minutes using a CytoFluor II fluorescence plate reader

### FRET VWF71 RBC Catalytic Assay

RBCs were isolated from wild-type and ADAMTS13-expressing C57BL/6 mice, washed, centrifuged, and resuspended in VWF71 assay buffer (50 mM HEPES, pH 7.4, 150 mM NaCl, 10 mM CaCl₂, 0.05% Tween- 20) at a concentration of 500,000 cells/µL. For each condition, 25 µL of the cell suspension was transferred to a 96-well plate, followed by the addition of 25 µL of 2.2 µM fluorescently labeled VWF71 substrate. Fluorescence kinetics were measured every 2.5 minutes over a 30-minute period using a PerkinElmer Victor fluorescence plate reader.

### FRET VWF73 Plasma Inhibition Assay

K562 cell lines engineered to express or lack ADAMTS13 were washed, centrifuged, and resuspended in 20 µL of plasma collected from patients with thrombotic thrombocytopenic purpura (TTP). Cells were incubated for 30 minutes at 4°C, followed by washing and resuspension in VWF73 assay buffer (Sensolyte). For each condition, 10 µL of the cell suspension (50,000 cells/µL) was dispensed into a 384- well plate, and 10 µL of fluorescently labeled VWF73 substrate (Sensolyte) was added. Fluorescence intensity was measured every 2.5 minutes for 30 minutes using a CytoFluor II fluorescence plate reader.

### vWF multimer analysis

Approximately 50,000 control or MTDCS-expressing K562 cells were washed twice with PBS and incubated in PBS buffer containing 2.2 μM of rhvWF for 1 hour at 37°C. Following incubation, cells were removed by centrifugation at 500 × g for 10 minutes. VWF multimer analysis was conducted on the supernatant using the Sebia Hydragel 11 von Willebrand Multimers kit (Sebia GmbH, Fulda, Germany) on the Sebia Hydrasis 2 platform, following the manufacturer’s instructions. Samples were diluted with sample diluent (Sebia), incubated at 45°C for 20 minute, and loaded onto a precast agarose gel (sample volume of 5 μL). Electrophoresis, in-gel immunofixation and VWF staining were done via manufacturer specifications. Gels were scanned and densitometry was performed using the Phoresis software version 6.1 (Sebia), as per manufacturer specifications.

### Flow cytometry

ADAMTS13 was detected using Invitrogen antibody MA5-24167 and either mouse IgG-PE (12-4010-82) or mouse IgG1-alexa-fluor 488 (A-11059). Enucleation was assessed by staining with Draq5 (following Thermo Fisher Scientific instructions) and FACS analysis.

### Blood counts

Samples were analyzed on an ADVIA 2120i automated hematology analyzer. Blood smears were made post inoculation and stained with Wright-Giemsa in order to detect the presence of schistocytes.

## Acknowledgments

The study was supported by grant NIH R01HL130764. We thank Kira Gritsman, Herb Lachman, Sanjeev Gupta and Henny Billett for useful discussions. Data in this paper are from a Thesis to be submitted in partial fulfillment of the requirements for the Degree of Doctor of Philosophy in the Biomedical Sciences, Albert Einstein College of Medicine.

## Authorship

K.S.R. designed research, performed research, analyzed data, and wrote the paper. S.Z. performed research and analyzed data. K.B. performed research and analyzed data (ADAMTS13-K562 cells). Z.Y. performed research and analyzed data. J.M. contributed vital new reagents (FRET-vWF71).

J.J.M. contributed analytical tools (vWF multimer analysis). E.O. performed research and analyzed data.

J.M.P. analyzed data (TTP model histology and pathology). S.R.C. contributed vital reagents (TTP plasma). E.E.B. designed research, analyzed data and wrote and approved the paper. Authors have no conflict of interest to declare.

