## Supplemental Data for "Proof-of-Concept in a Murine Model of Treatment of Thrombotic Thrombocytopenic Purpura Using Engineered Red Blood Cells"

**Supplemental Figure legends:**

**Figure S1: Fusion constructs tested at the AAVS1 locus.**

**A:** Schematic map of the fusion constructs inserted at the AAVS1 site, along with a representative PCR assay confirming successful integration at the AAVS1 locus.

**B:** Summary table detailing the different constructs tested in K562 cells at the AAVS1 locus, including variations in membrane anchors and ADAMTS13 fragments.

**Figure S2: Transfusion of AD5M-Reticulocytes.**

**A:** Approximately 300 million AD5 reticulocytes were transfused into control mice, and ADAMTS13 expression on the RBC membrane was monitored over time by FACS using an anti-ADAMTS13 antibody. Expression of ADAMTS13 disappeared over time.

**B:** Approximately 200 million AD5 reticulocytes were labeled with the lipophilic dye DID and transfused into control mice. ADAMTS13 surface expression was assessed by FACS using an anti-ADAMTS13 antibody. The results demonstrate that while the DID-labeled cells persist in circulation, ADAMTS13 membrane expression diminishes over time.

**Figure S3: Schematic illustrating the design of the proof-of-concept experiment.**

Created in BioRender. Roberts, K. (2025) https://BioRender.com/wk3uhzh

**Figure S4: Transfusions of AD5M-Reticulocytes prevent organ damage in the Schiviz TTP mouse model.**

Representative micrographs (4× magnification) of formalin-fixed, paraffin-embedded murine heart and kidney sections stained with H&E, obtained 4 days after transfusion with either control or AD5M-reticulocytes. Reticulocytes were transfused 3 hours prior to rhvWF injection to induce TTP symptoms. All mice (4/4) transfused with control reticulocytes showed cardiac leukocyte infiltration. In contrast, no infiltration was observed in mice receiving AD5M-reticulocytes. These results indicate that AD5M-reticulocyte transfusion prevents organ damage in this TTP model. Scale bar is 500 μm.

### knock-in at AAVS1

(Chr19: PPP1R12C)

A

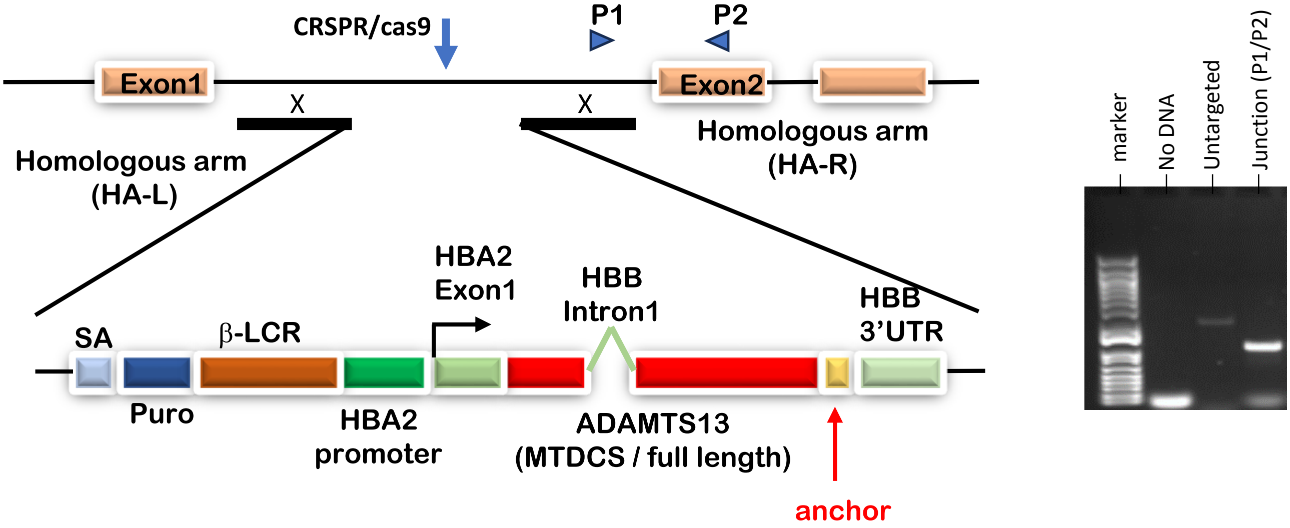


B

| Plasmid name | ADAMTS13 | Anchor | Mutations |
|---|---|---|---|
| **P843 (AD5 GPI)** | MTDCS | DAF GPI | WT |
| **P849 (AD5M GPI)** | MTDCS | DAF GPI | Gain-of function mutations |
| **P881 (AD5 GPB)** | MTDCS | GPB full-length | WT |
| **P882 (AD5 TM)** | MTDCS | GPB transmembrane domain | WT |
| **P883 (AD13 GPB)** | Full-length | GPB full-length | WT |
| **P884 (AD13 TM)** | Full-length | GPB TM | WT |
| **P892 (K608N AD5 GPB)** | MTDCS | GPB full-length | K608N |

**Figure S1**

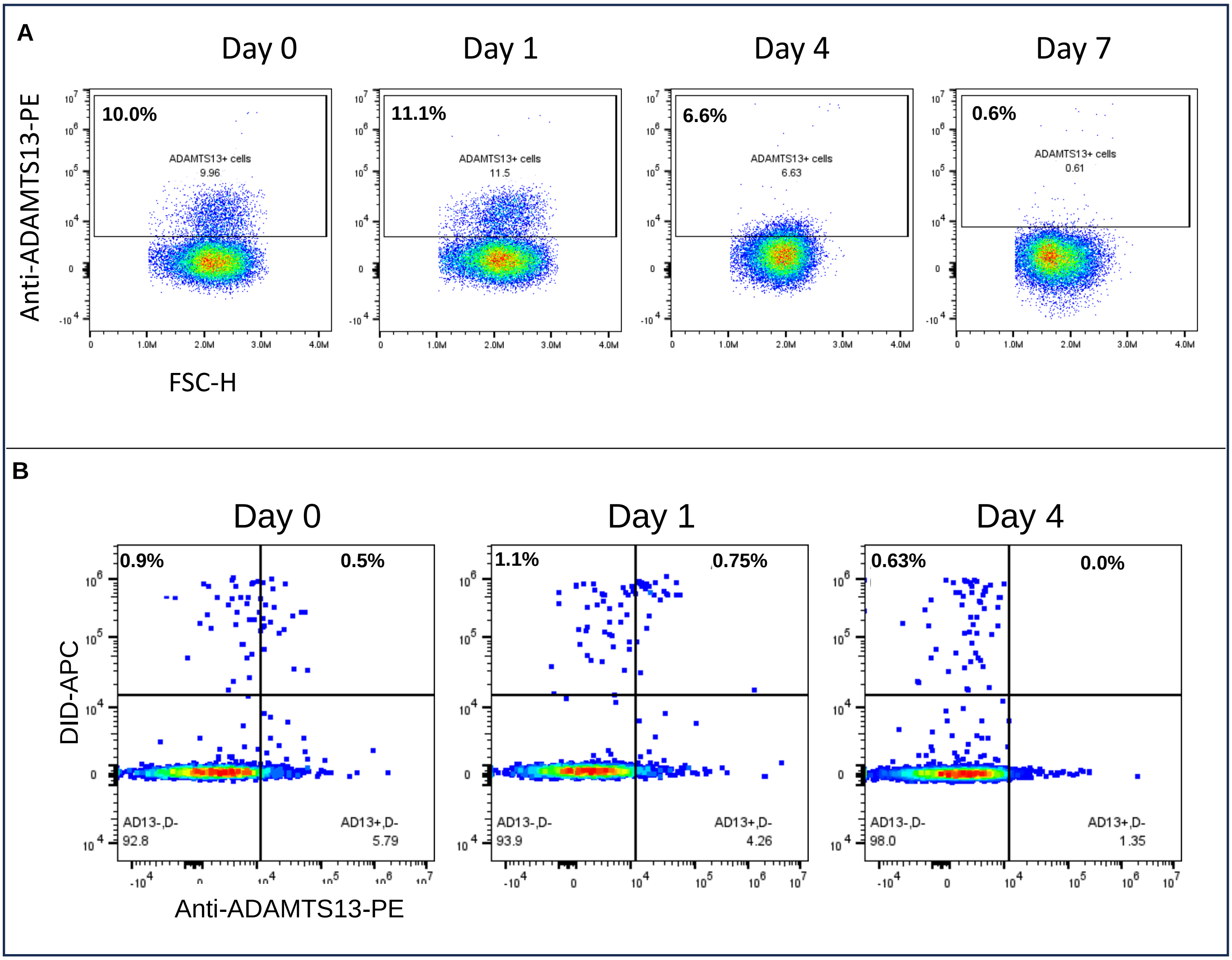
A
Day 0
Day 1
Day 4
Day 7
Anti-ADAMTS13-PE
10.0%
ADAMTS13+ cells
9.96
11.1%
ADAMTS13+ cells
11.5
6.6%
ADAMTS13+ cells
6.63
0.6%
ADAMTS13+ cells
0.61
FSC-H
B
Day 0
Day 1
Day 4
DID-APC
0.9%
0.5%
AD13-,D-
92.8
AD13+,D-
5.79
1.1%
0.75%
AD13-,D-
93.9
AD13+,D-
4.26
0.63%
0.0%
AD13-,D-
98.0
AD13+,D-
1.35
Anti-ADAMTS13-PE

**Figure S2**

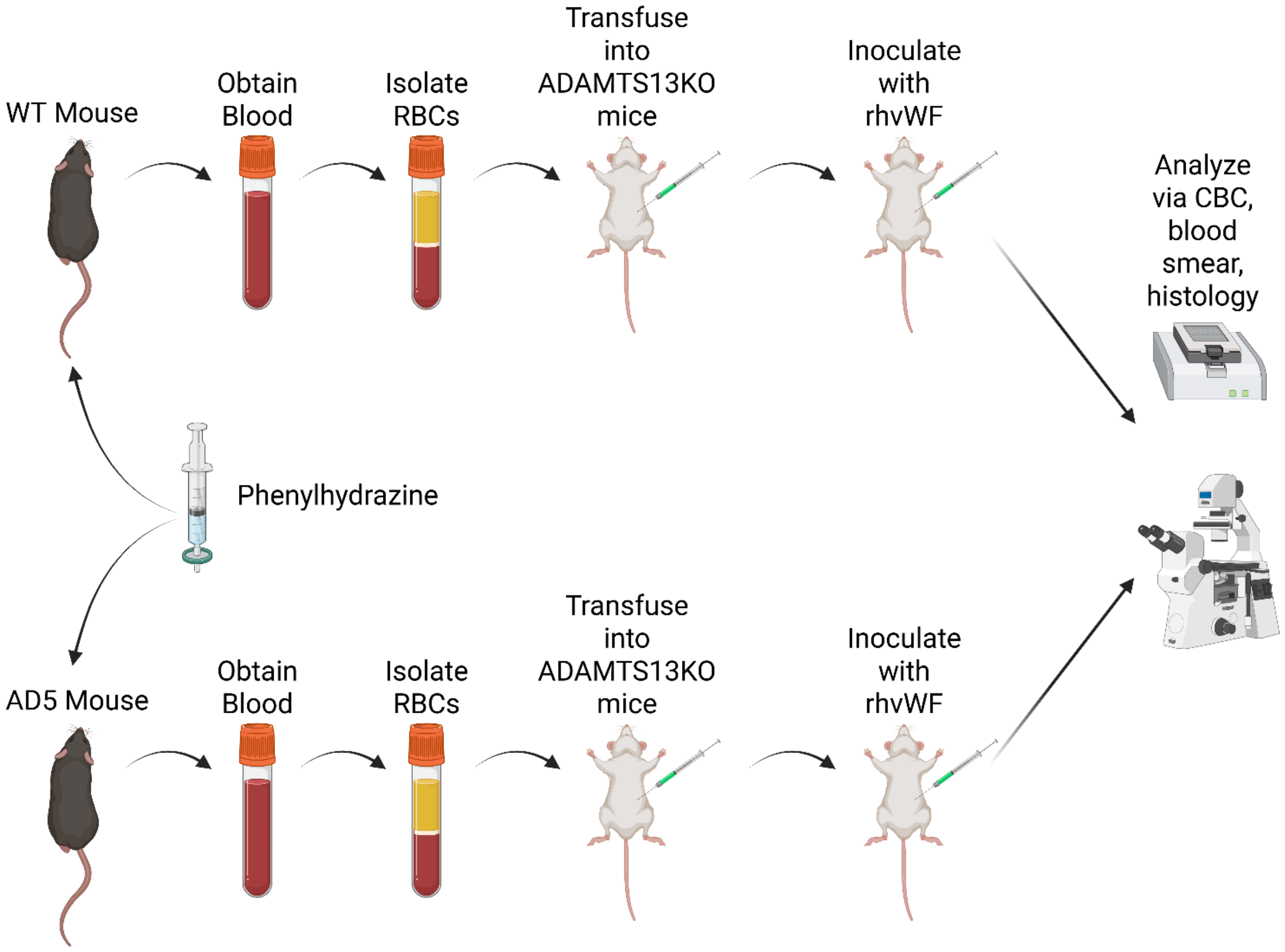
WT Mouse
Obtain Blood
Isolate RBCs
Transfuse into ADAMTS13KO mice
Inoculate with rhvWF
Analyze via CBC, blood smear, histology
Phenylhydrazine
AD5 Mouse
Obtain Blood
Isolate RBCs
Transfuse into ADAMTS13KO mice
Inoculate with rhvWF

**Figure S3**

Control Transfusion

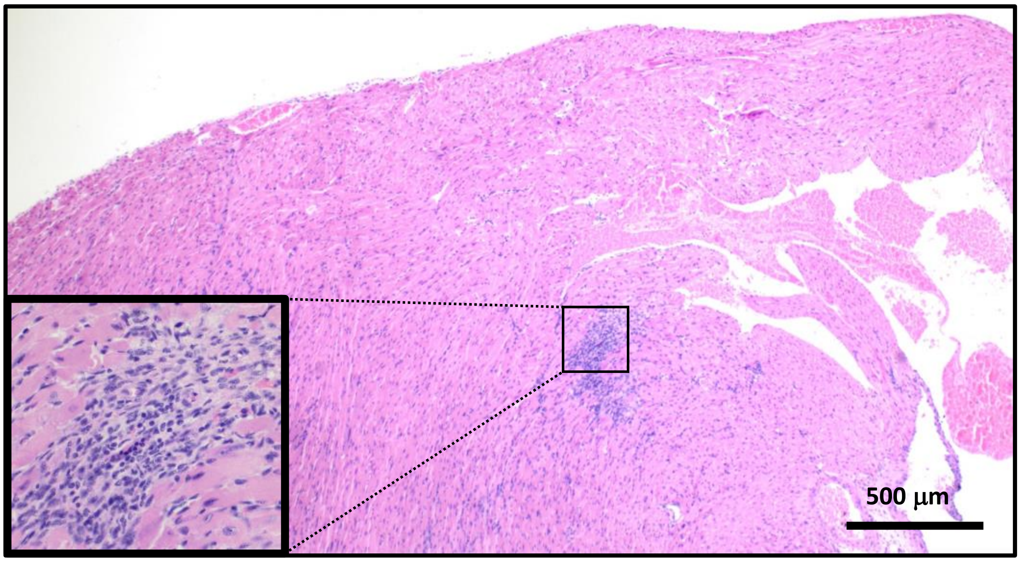


AD5M Transfusion

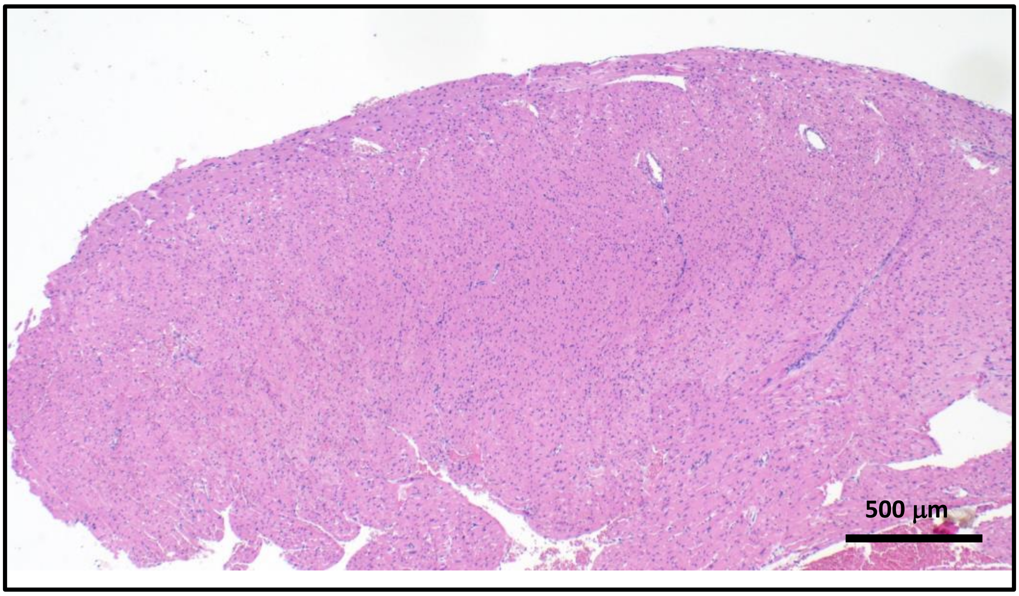


**Figure S4**
